# Spindle assembly checkpoint satisfaction occurs through end-on but not lateral kinetochore-microtubule interactions under tension

**DOI:** 10.1101/105460

**Authors:** Jonathan Kuhn, Sophie Dumont

**Affiliations:** Tetrad Graduate Program, University of California, San Francisco, San Francisco, CA, 94143; Department of Cell and Tissue Biology, University of California, San Francisco, San Francisco, CA, 94143; Department of Cell and Molecular Pharmacology, University of California, San Francisco, San Francisco, CA, 94143

## Abstract

To ensure accurate chromosome segregation, the spindle assembly checkpoint (SAC) prevents anaphase until all kinetochores attach to the spindle. What signals the SAC monitors remains unclear. We do not know the contributions of different microtubule attachment features, or tension from biorientation, to SAC satisfaction in normal mitosis - or how these possible cues change during attachment. Here, we quantify concurrent Mad1 intensity, reporting on SAC silencing, and real-time attachment geometry, occupancy, and tension at individual mammalian kinetochores. We show that Mad1 loss from the kinetochore occurs in switch-like events with robust kinetics, and that metaphase-like tension across sister kinetochores is established just before Mad1 loss events at the first sister. We demonstrate that CenpE-mediated lateral attachment of the second sister can persistently generate this metaphase-like tension prior to biorientation, likely stabilizing sister end-on attachment, yet cannot induce Mad1 loss from that kinetochore. Instead, Mad1 loss begins after several end-on microtubules attach. Thus, end-on attachment provides geometry-specific molecular cues, or force on specific kinetochore linkages, that other attachment geometries cannot provide.

**Summary:** The spindle assembly checkpoint (SAC) delays anaphase until kinetochores are properly attached to the spindle. The authors demonstrate that the SAC monitors geometry-specific molecular cues, or force on specific kinetochore linkages, that “end-on” but not “lateral” attachments generating persistent tension can provide.

## Introduction

The spindle assembly checkpoint (SAC) ensures correct partitioning of the genome (London and Biggins, 2014b; Musacchio and Salmon, 2007) by preventing the onset of anaphase until all sister kinetochores are attached to the spindle (Rieder et al., 1995). The level of SAC proteins at kinetochores regulates cell cycle progression (Collin et al., 2013; Dick and Gerlich, 2013; Heinrich et al., 2013). Specifically, the removal of Mad1 from attached kinetochores controls the anaphase-inhibitory signal (Maldonado and Kapoor, 2011). Both tension from biorientation (McIntosh, 1991) and microtubule attachment have been proposed to control Mad1 loss from kinetochores in mammalian cells (Etemad and Kops, 2016; Pinsky and Biggins, 2005). While tension across the centromere is not essential for SAC satisfaction (Rieder et al., 1995), tension within (and across) a single kinetochore may be (Maresca and Salmon, 2009; Uchida et al., 2009). Across which linkages tension could be monitored, and whether that tension would be necessary, sufficient, or neither, remains unclear (Etemad et al., 2015; Magidson et al., 2016; Smith et al., 2016; Tauchman et al., 2015). Instead, or in addition, the SAC may monitor the presence of microtubules or of a specific attachment feature such as whether the kinetochore binds to the end (end-on) or side (lateral) of microtubules (i.e. geometry), how many microtubules it binds (occupancy), and the timescale over which it remains attached to any or a given-microtubule (lifetime).

We do not know what physical changes ultimately trigger Mad1 loss and, critically, we do not know how changes in tension and attachment features map to those of Mad1 at individual kinetochores during mammalian spindle assembly. One barrier is that to this point we have only visualized tension, attachment and SAC signaling together in fixed cells. Challenges to live imaging these dynamics include concurrently visualizing individual kinetochores moving in 3D, dim microtubule structures, and dim Mad1 - and doing so at high resolution over long periods. Since tension and attachment candidate cues both evolve during mitosis (Magidson et al., 2011; McEwen et al., 1997), and often co-vary and go through short-lived intermediates, uncoupling their contributions has been difficult. In particular, it is not yet clear if lateral attachments, which utilize motors rather than Ndc80 for binding microtubules, are able to trigger SAC silencing (Cheeseman and Desai, 2008; Kops and Shah, 2012; Nezi and Musacchio, 2009), and whether the answer to this is based on the specificity of their molecular interface, or to a different stability or force-generating ability. Determining which kinetochore interfaces can and cannot trigger SAC silencing upon binding to, or being pulled on by, microtubules is essential to understanding what the kinetochore monitors to control cell cycle progression.

Here, we develop an approach to quantitatively map in real-time the structural dynamics of centromere tension, attachment geometry, and attachment occupancy onto those of Mad1 signaling at individual mammalian kinetochores during spindle assembly. Together, our work reveals the space-time trajectory of a sister pair to SAC silencing, and indicates that engagement of microtubule ends is the trigger for SAC silencing. We demonstrate that CenpE-based lateral attachments can generate long-lived force, and are thus well-suited to stabilize end-on attachments before biorientation, but they cannot satisfy the SAC. Thus, end-on attachment must provide specific molecular cues - or force on a specific linkage - to control the SAC that other persistent, force-generating attachments cannot provide.

## Results and discussion

To measure the real-time kinetics of Mad1 depletion once initiated at individual kinetochores, we used a two-color reporter consisting of EYFP-Mad1 and CenpC-mCherry (Fig. 1A). Mad1 localization is necessary and sufficient for SAC activation (Maldonado and Kapoor, 2011), its N-or C-terminal EYFP fusions behave similarly (Shah et al., 2004), and Mad1 binding and dissociation kinetics are well understood on unattached - but not attached - kinetochores (Dick and Gerlich, 2013; Howell et al., 2004; Shah et al., 2004). In turn, CenpC is a stable kinetochore component (Shah et al., 2004). The ratio of Mad1 to CenpC intensities controls for variation in kinetochore size and for out of focal plane movements. We imaged this reporter live over four focal planes with ~10 s resolution in mammalian PtK2 cells. These cells are ideal to track kinetochores and map physical attachment and tension changes as they are large, flat, and have few chromosomes. We tracked individual kinetochores from prophase or prometaphase to metaphase using CenpC-mCherry, and quantified reporter intensities (Fig. 1A).

**Figure 1:**
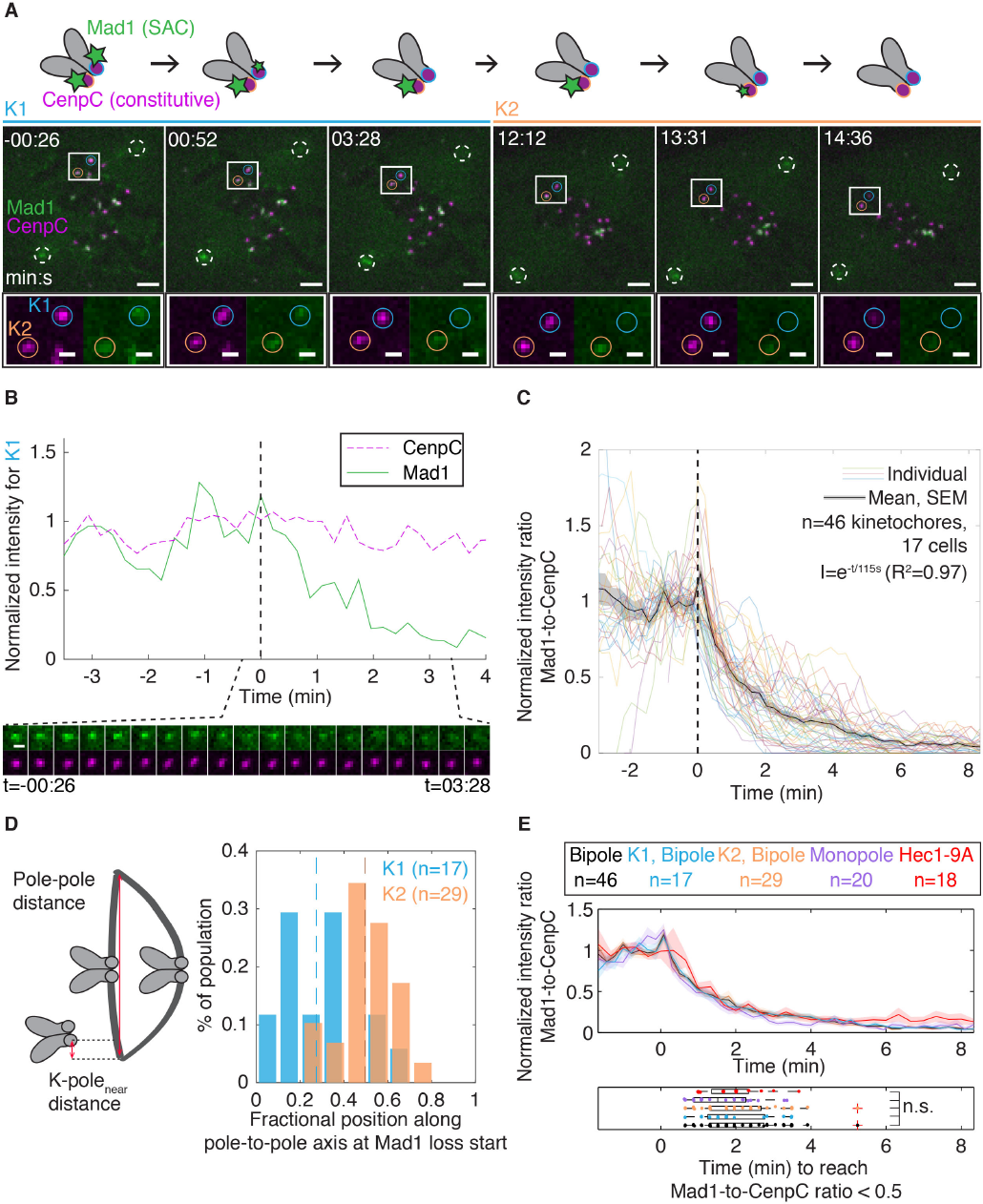
Mad1 loss at individual mammalian kinetochores is a switch-like process with robust, stereotypical single exponential kinetics. Timelapse imaging (maximum intensity projection) of representative SAC satisfaction kinetics (EYFP-Mad1, green) at individual kinetochores (CenpC-mCherry, magenta) during spindle assembly in a PtK2 cell. Full circles identify first (K1, blue) and second (K2, orange) sisters in a pair to lose Mad1, and white dashed circles identify spindle poles. Scale bars=3 μm (large) and 1 μm (zoom), and t=0 indicates the start of Mad1 loss on K1. See also Video 1. **(B)** Mad1 (solid green) and CenpC (dashed magenta) intensities for K1 in (A) around SAC satisfaction. Timelapse (bottom) of K1 at 13 s intervals. Scale bar=1 μm. **(C)** Individual traces, mean and SEM of the Mad1-to-CenpC intensity ratio (I) over time (t) around SAC satisfaction, with traces synchronized at t=0. Mad1 loss kinetics are switch-like and stereotypical. **(D)** Distribution of the fractional position along the pole-to-pole axis ((kpole_near_)/( pole-pole)) where kinetochores start to lose Mad1. Dashed lines indicate the mean position for each sister. The first (K1) and second (K2) sister in a pair to lose Mad1 satisfy the SAC at different spindle positions (p<0.01): K1 close to its pole, and K2 near the metaphase plate. **(E)** Mean and SEM (top) of the Mad1-to-CenpC intensity ratio around SAC satisfaction (with t=0 Mad1 loss start), and distribution (bottom) of times to reach a threshold intensity ratio, in bipolar cells (black, all kinetochores), different kinetochores (K1 and K2) in these cells (blue and orange),in monopolar spindles (STLC-treated; Video 2) which have lower tension (purple), and Hec1-9A-expressing cells which have higher tension and attachment levels (red). In all cases, Mad1 loss kinetics are indistinguishable from control (n.s. is for p>0.05).

We found that Mad1 levels at individual kinetochores are stable during spindle assembly (Mad1-ON state), until they drop sharply to background (Mad1-OFF state) while CenpC levels stay constant (Fig. 1A, B, Video 1). We did not find intermediate steady-states with intensities between those of Mad1-ON and-OFF states, and did not observe any significant re-recruitment of Mad1 upon the completion of Mad1 loss. To determine the distribution of Mad1 loss rates, we aligned all Mad1 loss events in time at the start of Mad1 loss (t=0). This revealed that Mad1 loss kinetics are switch-like and strikingly similar over kinetochores and cells (n=46 kinetochores in 17 cells; Fig. 1C). The kinetics of Mad1 loss are well fit by a single exponential (t1/2=79 s, R^2^=0.97; Fig. 1C), with only a marginal fit improvement with a double exponential (R^2^=0.99). This suggests that the process of removing Mad1 from kinetochores has one rate-limiting step. This single event could, for example, be the turnover of phosphorylation marks involved in recruiting Mad1 (Nijenhuis et al., 2014), or Mad1 removal by dynein (Howell et al., 2001).

To probe the events that govern Mad1 loss, we examined Mad1 removal at individual kinetochore pairs and in different conditions. Mad1 loss events at each sister were broadly distributed in time (10.6±9.5 min apart; Fig. S1A), and thus pairs with a single SAC-satisfying sister exist (Gorbsky and Ricketts, 1993), and were broadly distributed in in space (Fig. 1D, S1B). The first sister in a pair to satisfy the SAC began losing Mad1 close to its spindle pole, often as it transitioned from poleward to away-from-pole movement; meanwhile, the second sister began losing Mad1 at a different location (p=0.0009), close to the metaphase plate, and typically after a sharp movement towards the plate that also aligned (Magidson et al., 2015) sisters along the pole-to-pole axis (Fig. S1C). Despite these differences, Mad1 loss events at the first and second kinetochores had indistinguishable kinetics after the onset of Mad1 loss (Fig. 1E). We then asked what physical properties of the kinetochore, if any, regulated this rate-limiting step in Mad1 loss. We hypothesized that tension or attachment, both proposed to be required for Mad1 loss, tune kinetics of Mad1 removal after its onset. To test this hypothesis, we imaged Mad1 in cells with monopolar spindles (Fig. S2A) which have lower tension (Fig. S2C), and in cells expressing Hec1-9A-mRuby2 (Fig. S2B) which have higher tension (Fig. S2C) and microtubule occupancy (Zaytsev et al., 2014). We use centromere stretch (interkinetochore (K-K) distance) as a reporter of tension: while centromere stretch is not itself necessary for SAC satisfaction, changes in centromere stretch imply changes in load on the k-fiber and across at least some kinetochore linkages. Despite these tension differences, Mad1 loss kinetics remained unchanged in monopoles (p=0.15, n=20 kinetochores, Fig. 1E; Video 2) and Hec1-9A cells (p=0.95, n=18 kinetochores; Fig. 1E). This suggests that after the onset of Mad1 loss, the rate of loss is insensitive to tension and attachment occupancy levels; once the SAC satisfaction decision is made, the kinetochore silences the SAC in a stereotypical event.

To gain insight into what events initiate Mad1 loss, we quantified how tension and attachment change before and around these stereotypical Mad1 loss events. We began by measuring the K-K distance of a sister pair before and during Mad1 loss (Fig. 2A-C, Video 3). In all cases mapped, the K-K distance of a single chromosome increased just before the first sister lost Mad1 (Fig. 2B-C): it started at 1.06 ± 0.06 µm for t<-2 min, indistinguishable (p=0.46) from that in nocodazole (n=10 pairs), and increased (p=0.008) to 2.14±0.32 µm by t=0, the start of Mad1 loss (Fig. 2D). The K-K distance as Mad1leaves the first kinetochore is higher in bipoles than in monopoles (2.14±0.32 µm versus 1.20±0.03, p=0.02; Fig. 2D and S2C), suggesting that an opposing force other than polar ejection force acts prior to silencing the first kinetochore in normal, bipolar mammalian mitosis. The K-K distance increase just before Mad1 loss, in all cases measured (Fig. 2E), is consistent with - but does not imply - tension across the kinetochore being necessary to initiate Mad1 loss. However, kinetochore pairs persist at a metaphase-level of tension (p=0.55) for minutes without Mad1 loss at the second sister (1.89±0.19 µm for t<-2 min, n=19 pairs; Fig. 2D) with no significant tension increase (p=0.96) at t=0 (1.89±0.14 µm). Thus, neither microtubule attachment nor the transmission of force (i.e. load-bearing) across the kinetochore, from chromosome to microtubule, is sufficient to satisfy the SAC in normal dividing cells.

**Figure 2:**
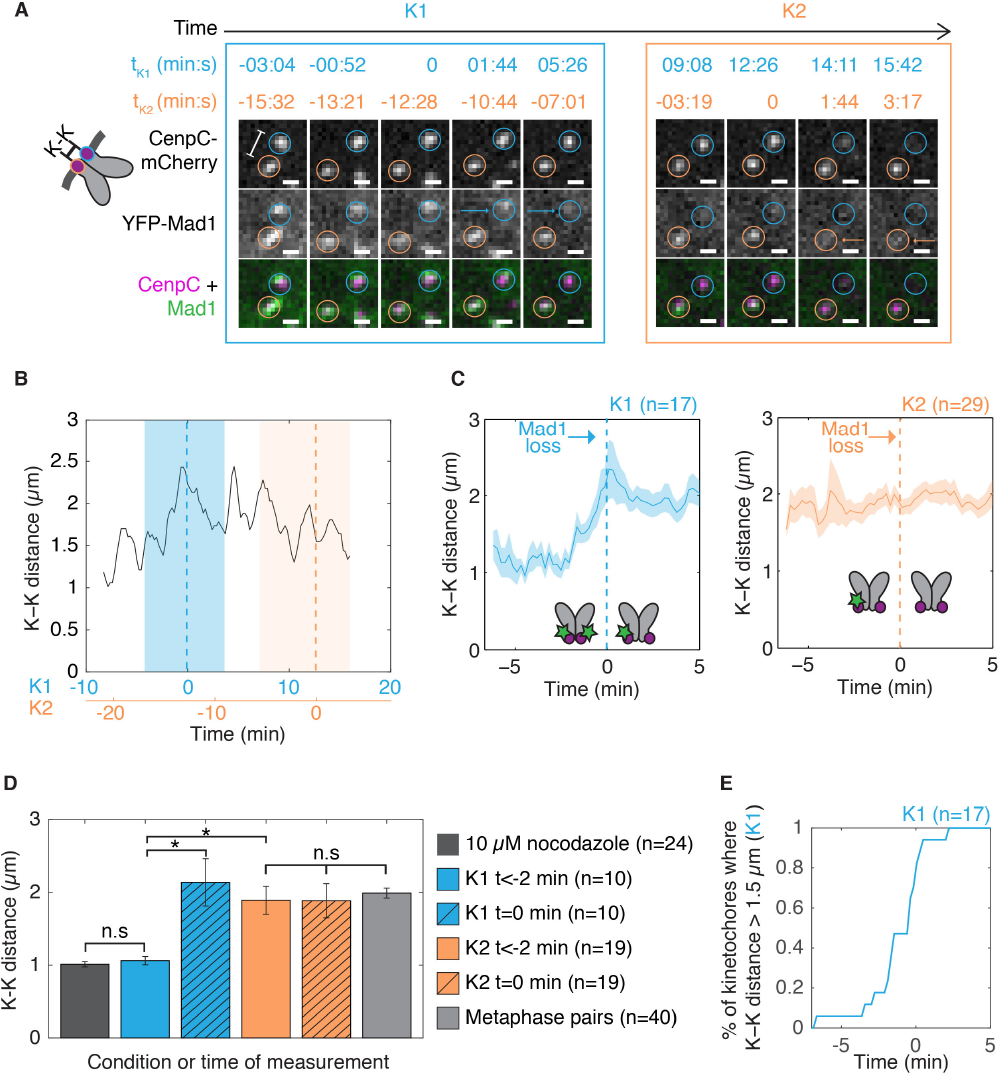
Kinetochores are under metaphase-level tension prior to Mad1 loss, but tension across the kinetochore is insufficient to initiate Mad1 loss. **(A)** Timelapse imaging (maximum intensity projection) of SAC satisfaction kinetics (EYFP-Mad1) concurrently with K-K distance (CenpC-mCherry) showing representative changes in centromere tension relative to Mad1 loss (start at t=0, followed by arrows) on K1 (blue) and K2 (orange) from PtK2 cell in Fig. 1A. See also Video 3. **(B)** K-K distance over time for the pair in (A), with smoothing over a 3-timepoint window, and t=0 (dashed lines) indicating the start of Mad1 loss for K1 (blue) and K2 (orange). **(C)** Mean and SEM of K-K distance over time plotted relative to Mad1 loss start (t=0) for each of K1 (left) and K2 (right). As in (A), tension increases prior to Mad1 loss at K1, but remains high for many minutes without loss of Mad1 at K2. **(D)** K-K distance at all timepoints before t=-2 min for K1 and K2, at t=0 for K1 and K2, and at reference points in separate experiments (metaphase kinetochores and 10 μM nocodazole). Measurements for K2’s t=-2 min do not include any data before K1’s t=0. Before Mad1 loss for K1, tension is low; in contrast (* p<0.01), pairs are already under high tension levels prior to Mad1 loss on K2. **(E)** Fraction of pairs with a K-K distance crossing>1.5 μm as time evolves relative to Mad1 loss start at K1 (t=0).

To uncover how kinetochores form persistent force-generating attachments without satisfying the SAC, we concurrently quantified Mad1 intensity and the position and intensity of microtubules at kinetochores in live cells. We expressed EYFP-Mad1 and mCherry-tubulin (Fig. 3A, Video 4). On pairs with one Mad1-ON sister (K2, orange), the Mad1-ON sister was associated with a microtubule bundle whose intensity continued past the kinetochore rather than terminating at it (Fig. 3A, Video 4). This is consistent with motor-driven lateral kinetochore-microtubule attachments, which we identified as cases where i) the tubulin intensity is equal on both sides of a kinetochore (Fig. 3B) and where ii) there is centromere tension (Kapoor et al., 2006) confirming productive microtubule engagement. Despite interactions with lateral microtubules (Fig. 3A-C) and high tension for long periods (Fig. 3D), the levels of Mad1 on these kinetochores did not change (n=8 kinetochores; Fig. 3E). Consistent with these attachments being powered by CenpE, a plus-end-directed kinesin, rigor inhibition of this motor with GSK-923295 (Wood et al., 2010) (Fig. 3F) generated kinetochore pairs stuck near a pole that failed to congress (Fig. 3G). These pairs remained with one laterally attached, Mad1-positive sister (Magidson et al., 2015) (n=10 kinetochores; Fig. 3H, Video 5), under some, albeit reduced, centromere tension (Fig. 3I). Thus, stable lateral microtubule-kinetochore attachments that generate long-lived metaphase centromere tension levels exist during normal mitosis, are mediated by CenpE, and are not sufficient to (even partially) satisfy the SAC. If the SAC monitors tension across the kinetochore, it must do so across a kinetochore linkage that is not put under sufficient load in these lateral attachments.

**Figure 3:**
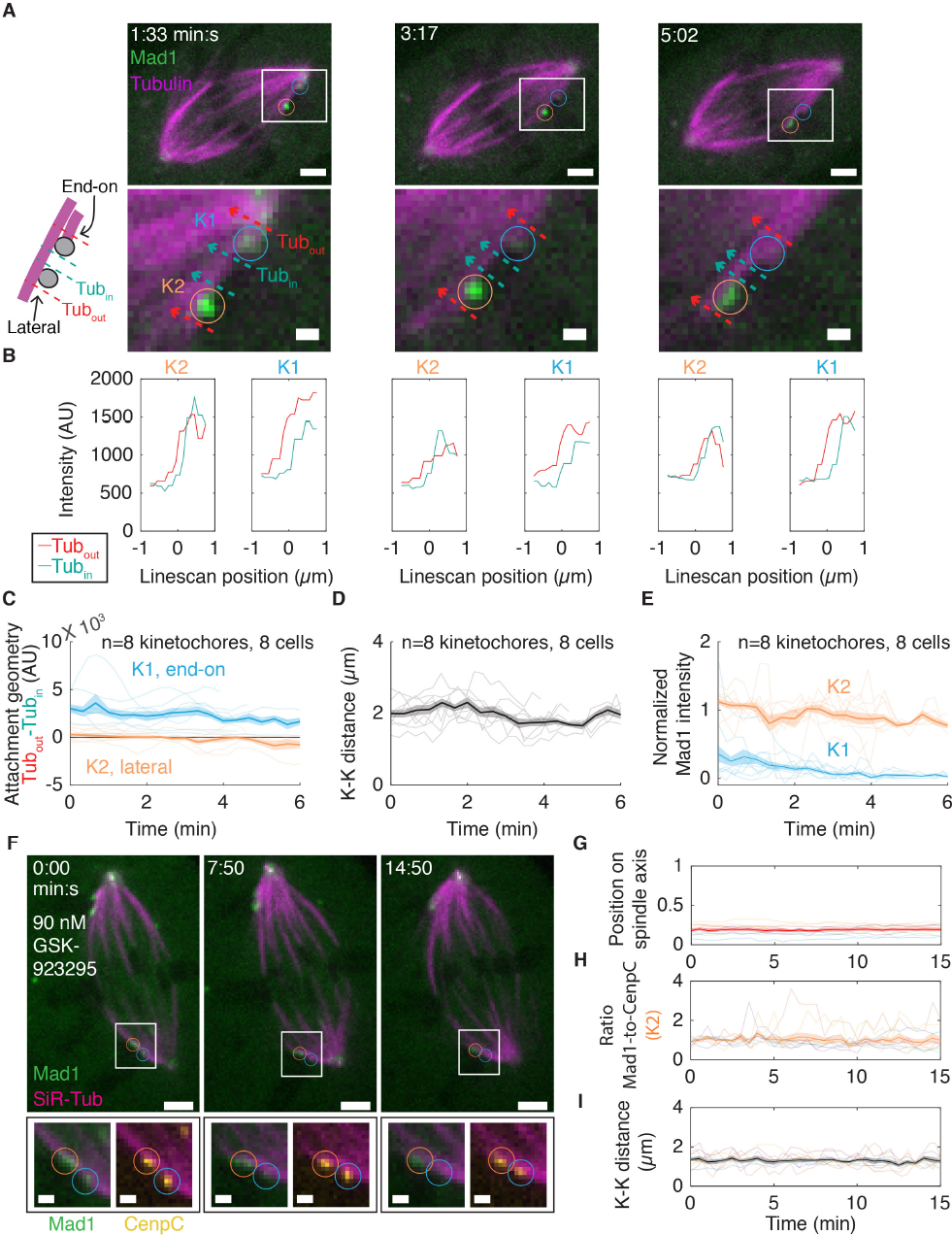
Lateral attachments generating long-lived metaphase-level centromere tension do not satisfy the SAC. **(A)** Timelapse imaging (maximum intensity projection) of representative SAC satisfaction kinetics (EYFP-Mad1) and microtubule attachment (mCherry-Tubulin) in a PtK2 cell. K2 (orange circle) is under tension and sits along the side of a neighboring kinetochore-fiber, suggesting a lateral attachment, but remains Mad1-positive while its sister K1 (blue circle) loses Mad1. Bottom images display the analysis depicted in (B). Scale bars=3 μm (large) and 1 μm (zoom), and t=0 indicates movie start. See also Video 4. **(B)** Analysis of microtubule geometry comparing the tubulin intensity (integrated linescans) inside (Tub_in_) and outside (Tub_out_) the pair in (A). The intensity difference is high on K1, indicating an end-on attachment, and is near zero on K2, indicating a lateral attachment. **(C)** Individual, mean and SEM of Tub_out_-Tub_in_ for K1 (blue) and K2 (orange) in pairs where K2 begins Mad1-positive. K1 and K2 have stable end-on and lateral attachments, respectively. **(D)** Individual, mean and SEM of K-K distances for the traces in (C), showing sustained metaphase-level centromere tension. **(E)** Individual, mean and Mad1 intensity for K1 (end-on, blue) and K2 (lateral, orange) in (C-D). Long-lived force-generating (D) lateral attachments (C) do not satisfy the SAC (E). **(F)** Timelapse imaging (maximum intensity projection) of representative SAC inactivation kinetics (EYFP-Mad1) and microtubule attachment (SiR-Tubulin) at a kinetochore pair (CenpC-mCherry) in a PtK2 cell where some kinetochores are locked in a CenpE-mediated lateral attachment (90nM GSK-923295 CenpE inhibitor) for >15 min. During this time, the pole-distal kinetochore (K2, orange circle) remains along the side of a microtubule with stable Mad1 levels. Scale bars=3 μm (large) and 1 μm (zoom), t=0 indicates movie start. See also Video 5. **(G-H)** Analysis (individual, mean and SEM) of **(G)** fractional position along the pole-to-pole axis (see Fig. 1D), **(H)** Mad1-to-CenpC intensity ratio and **(I)** K-K distance for such K2 kinetochores highlighted in (F), where t=0 indicates movie start. Mad1 levels are stable (H) at these kinetochores that remain stuck away from the metaphase plate (G) and general some, but not full, tension (I). These observations are consistent with CenpE powering the above (A-E) long-lived forcegenerating lateral attachments.

Finally, to probe how microtubule attachment geometry and occupancy changed before and around Mad1 loss, we imaged three-color cells expressing EYFP-Mad1 and CenpC-mCherry, and stained with the far-red microtubule dye SiR-Tubulin (Lukinavicius et al., 2014) (Fig. 4A, Video 6). We captured Mad1 loss events for the second kinetochore in a pair, and concurrently quantified (Fig. 4A, B) the dynamics of Mad1 intensity (Fig. 4C), microtubule attachment geometry and occupancy (Fig. 4D) and centromere tension (Fig. 4E) at single kinetochores. Strikingly, signature Mad1 loss events (Fig. 4C) always coincided with a sharp increase in end-on microtubule occupancy levels (n=21 kinetochores; Fig. 4B, D). As the first several microtubule ends bind from t=-100 s to t=0 (p=0.003; Fig. 4D), there is no corresponding decrease in Mad1 levels (Fig. 4C). After several end-on microtubules have bound, which we estimate to be half a mature kinetochore-fiber (thus about 10-12 microtubules in PtK cells (McEwen et al.,1997)), Mad1 loss then initiates (t=0), before end-on attachment levels reach their mature kinetochore-fiber levels at around t=100 s (p=0.001). Over this same time period, there was no change in centromere tension on these kinetochores (Fig. 4E). Together, our work indicates that the trigger for Mad1 loss is kinetochore engagement to microtubule ends, and that this engagement provides a unique, geometry-specific cue - whether molecular or physical - that other persistent force-generating attachments cannot provide.

**Figure 4:**
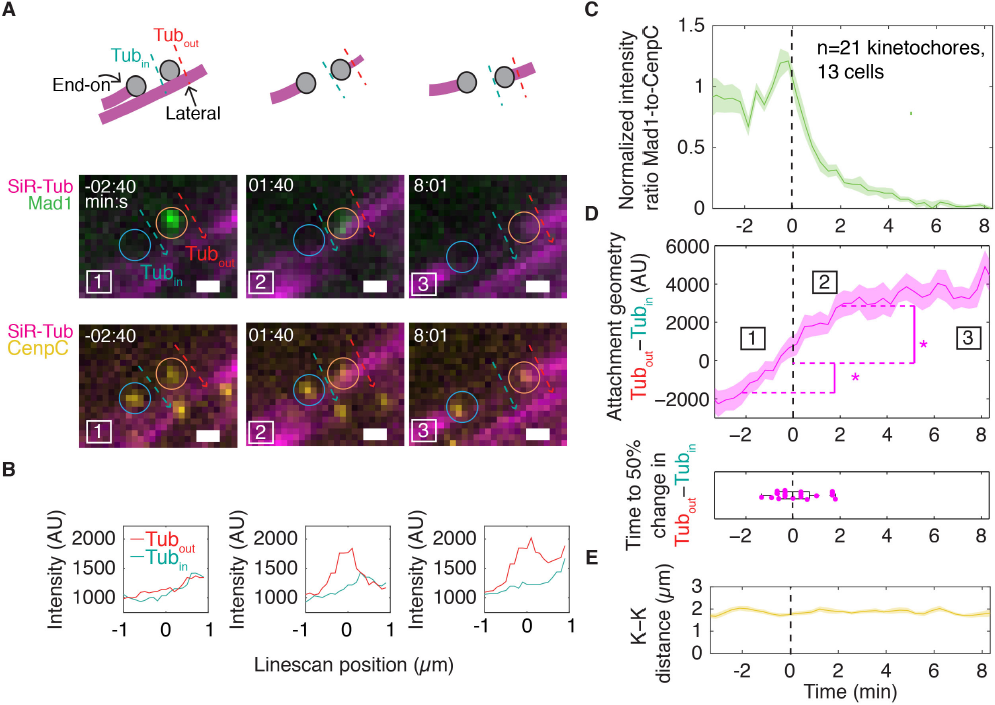
Mad1 loss begins rapidly after end-on attachment initiation and before a full kinetochore-fiber forms. **(A)** Timelapse imaging (maximum intensity projection) of a representative kinetochore pair’s SAC satisfaction kinetics (EYFP-Mad1; Mad1 loss start at t=0), attached microtubules’ geometry and intensity (SiR-Tubulin) and centromere tension (CenpC-mCherry) in a PtK2 cell. Dotted lines illustrate analysis shown in (B). Scale bars=1 μm. See also Video 6. **(B)** Microtubule attachment geometry (and occupancy) analysis showing an increase in Tub_out_- Tub_in_ as an end-on attachment forms, corresponding to images in (A). A negative value indicates that the kinetochore is near the end of its lateral microtubule track. **(C-E)** Concurrent quantification (mean, SEM) of **(C)** Mad1-to-CenpC intensity ratio, **(D)** microtubule attachment geometry (Tub_out_- Tub_in_), and **(E)** tension (K-K distance) around SAC satisfaction, with t=0 indicating the start of Mad1 loss on K2. Boxed numbers map to images in (A). Mad1 loss starts rapidly after end-on attachment initiation, when less than (* p<0.01, pairwise t-test) a full complement of microtubules are bound; about half of a full complement around t=0 (D, bottom). Meanwhile, there is no significant change in K-K distance (E).

Many elegant studies have used genetic and chemical perturbations to change tension and kinetochore-microtubule attachments, and thereby probe the events triggering Mad1 depletion. Here, our approach was to image naturally occurring centromere tension and attachment changes during spindle assembly - and to map those to the real-time kinetochore SAC silencing response. This allowed us to map a trajectory of events leading to SAC satisfaction in unperturbed cells (Fig. 5). Key to this work, we identified discrete signature events where Mad1 leaves individual kinetochores with robust, stereotypical kinetics. In the trajectory we map, likely one of a few, the first sister loses Mad1 close to its pole, after the second sister laterally attaches and generates tension through CenpE. This lateral attachment generates long-lived metaphase-like tension, well-suited to stabilize the first sister’s end-on attachment while both sisters are still connected to the same pole, but it cannot induce Mad1 loss on its kinetochore. Instead, this second sister loses Mad1 near the metaphase plate, after binding several end-on microtubules from the correct pole.

**Figure 5:**
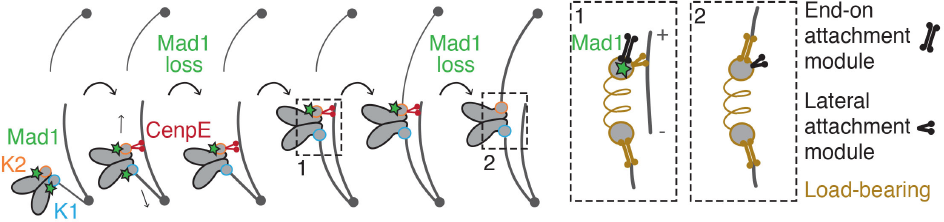
Model for attachment trajectory and triggering cue leading to SAC silencing of sister kinetochores. The first kinetochore (K1, blue circle) loses Mad1 (green star) and satisfies the SAC near its pole, after its sister (K2, orange circle) attaches to microtubules and generates force. In the trajectory we map, K2 laterally attaches through CenpE (red motor), generating tension that can stabilize end-on attachment of K1 - helping K1 bypass tension-based inhibition of initial end-on attachments. Despite being able to transmit force and bear load from the outer to inner kinetochore (Box 1, gold), this CenpE-based lateral attachment does not induce Mad1 loss at K2. CenpE pulls the pair towards the metaphase plate, where K2 forms end-on microtubule attachments to its pole and rapidly loses Mad1. SAC satisfaction must be triggered by a geometry-specific cue unique to end-on attachment that CenpE-based attachments (even persistent force-generating ones) cannot supply. This cue could, for example, be binding interactions specific to end-on attachments, or deformation of a linkage that only bears sufficient load in an end-on attachment (Box 2, gold).

The contributions of tension and attachment (lifetime, geometry, occupancy) have been hard to decouple. Here, we identify a long-lived state with high centromere tension and no detectable end-on attachment, revealing that neither the presence of long-lived tension somewhere in the kinetochore nor persistent microtubule attachment is sufficient to initiate Mad1 loss. Consistently, Mad1 had been found at some laterally-attached kinetochores in fixed cells (Dick and Gerlich, 2013; Drpic et al., 2015; Magidson et al., 2015; Shrestha and Draviam, 2013); however, whether the captured attachments were stable and generated sustained force - and therefore if end-on geometry was the only missing element of a correct attachment - was not known. While lateral attachments persist in mature metaphase plates (Magidson et al., 2011), may allow anaphase entry in some scenarios (O'Connell et al., 2008) and have been proposed to help SAC protein stripping (Howell et al., 2000) in mammalian cells, end-on attachment is necessary for SAC satisfaction in normal mitosis. What are the minimal events sufficient for SAC satisfaction? We find that stereotypical Mad1 loss begins without a full complement of microtubules; further, the different kinetics of k-fiber formation and Mad1 loss suggest that there is not simply a linear relationship between them. Mapping the precise relationship between microtubule occupancy and SAC signaling will require tools to disrupt the rapid k-fiber formation (McEwen et al., 1997) we observe, as high as 5-8 microtubules/min.

While our work shows that the presence of tension across kinetochore linkages, from chromosome to microtubule, in not sufficient for SAC satisfaction, in unperturbed mitosis we never observed Mad1 loss prior to both sisters attaching and generating tension. Our data strongly suggests that lateral attachments, which may be differently sensitive (Kalantzaki et al., 2015) to destabilization at low tension, facilitate end-on attachment formation and maintenance by generating tension (Foley and Kapoor, 2013; Khodjakov and Pines, 2010; King and Nicklas, 2000; Nezi and Musacchio, 2009; Nicklas et al., 2001) at the kinetochore-microtubule interface prior to biorientation. While polar ejection forces may generate tension across these kinetochores (Cane et al., 2013; Drpic et al., 2015), waiting for tension associated with bidirectional force before beginning SAC silencing - as observed in all events we monitored - could ensure that the pair is on a path to correct attachment before SAC satisfaction of the first sister. Methods to decouple attachment and tension across the kinetochore will be needed to determine if the latter is necessary for SAC satisfaction. Critically, we note that the inability of persistent force-generating lateral attachments to satisfy the SAC ensures that only bioriented attachments can both stabilize kinetochore-microtubule interactions and satisfy the SAC to control cell cycle progression. The imaging approach we developed, employed in different molecular backgrounds and with different fluorescent SAC reporters, should help uncover the cascade of events linking end-on microtubule engagement to SAC satisfaction.

The selectivity of the SAC for end-on attachments raises the question of what mechanism confers plus-end specificity. For example, the plus-end may engage a geometry-specific kinetochore interface because of its unique structure and dynamics, or plus-end geometry may allow kinetochore components to engage with more microtubules. One potential mechanism is that Mps1, a kinase upstream of Mad1 localization (London and Biggins, 2014a), is regulated in an end-on specific manner. Lateral attachments through CenpE may not compete with Mps1 for Ndc80 binding (Hiruma et al., 2015; Ji et al., 2015), or may not access the proper kinetochore interface to pull Mps1 away from its substrates (Aravamudhan et al., 2015). Lateral attachments satisfy the SAC in budding yeast (Krefman et al., 2015; Shimogawa et al., 2010), and the pathways that control dynein-dependent SAC protein stripping may also help confer end-on geometric specificity (Gassmann et al., 2010; Matson and Stukenberg, 2014) in mammals. If tension-based deformations within a kinetochore are important for SAC signaling, our findings suggest that these deformations must be highly specific to end-on microtubule attachments. Further, our work suggests that if tension is sensed, it would likely be sensed at a linkage outside of the junction where CenpE-and Ndc80-based attachments both transmit force (i.e. bear load) to the centromere (Fig. 5, Boxes 1-2). Force on different molecular interfaces at the kinetochore (Wynne and Funabiki, 2015), supporting either lateral or end-on binding, may differently propagate through the kinetochore and differently regulate kinetochore structure and SAC signaling (Fig. 5). Looking forward, it will be important to determine what kinetochore structural and biochemical changes take place when lateral and end-on attachments form.

## Materials and methods

**Cell culture and transfection:** PtK2 EYFP-Mad1 cells (Shah et al., 2004) (gift from Jagesh Shah, Harvard Medical School) were cultured as previously described (Elting et al., 2014). For imaging, cells were plated on 35 mm dishes with #1.5 poly-d-lysine coated coverslips (MatTek) and media was switched to identical media without phenol red 24 h prior to imaging. Cells were transfected using Fugene6 or Viafect (Promega) and imaged 36-48 h after transfection with mCherry-α-tubulin (gift from Michael Davidson, Florida State University), mCherry-CenpC (gift from Aaron Straight, Stanford University), or Hec1-9A-mRuby2 (mRuby2 (gift from Michael Davidson) was swapped for EGFP in Hec1-9A-EGFP (Guimaraes et al., 2008) (gift from Jennifer DeLuca, Colorado State University)).

**Drug and dye treatments:** To make monopolar spindles, 5 μM STLC (Sigma-Aldrich) was added 15 min before imaging (10 mM stock). To rigor CenpE to microtubules, 90 nM GSK-923295 (MedChem Express) was added 15 minutes before imaging (30 µM stock) (Magidson et al., 2015). To visualize tubulin as a third color, 100 nM SiR-Tubulin dye (Cytoskeleton, Inc.) was added 1 h prior to imaging (1 mM stock), along with 10 µM verapamil (Cytoskeleton, Inc.; 10 mM stock) to prevent dye efflux.

**Imaging:** Live imaging was performed on an inverted (Eclipse Ti-E; Nikon), spinning disk confocal (CSU-X1; Yokogawa Electric Corporation) microscope as previously described (Elting et al., 2014) for two-color imaging. For three-color imaging, a Di01-T405/488/568/647 head dichroic (Semrock) was used, along with a 642 nm (100 mW) diode laser and an ET690/50M emission filter (Chroma Technology Corp). Cells were imaged by phase contrast (200-400 ms exposures) and fluorescence (40-75ms exposures) in four z-planes spaced 350 nm apart every 13-30 s, with a 100× 1.45 Ph3 oil objective through a 1.5× lens (Metamorph 7.7.8.0; Molecular Devices). All images were collected at bin = 2 (to improve imaging contrast for dim Mad1 and microtubule structures), 5x pre-amplifier gain, and no EM gain (210 nm/pixel). Cells were imaged at 30 °C, 5% CO2 in a closed, humidity-controlled Tokai Hit PLAM chamber. The only image processing done prior to display were maximum intensity projections at each timepoint and (for videos only) linear scaling up of the image size in ImageJ.

## Data Analysis

**Tracking:** Kinetochore pairs were visually identified by coordinated motion and selected for analysis if they stayed away from other kinetochores and if at least one sister lost Mad1 during the movie. All further analysis was done within Matlab (Mathworks). If mCherry-CenpC was present, kinetochores were tracked using SpeckleTracker (Wan et al., 2012); if it was not present (Fig. 3), kinetochore tracking was done manually using Mad1, tubulin, and phase contrast images and custom software. Spindle poles were tracked manually.

**Intensity measurements:** To measure EYFP-Mad1 and mCherry-CenpC intensities at each time point, movies were thresholded by setting to zero all pixels <2 standard deviations above image background at the first frame. For each time point, the intensities of all pixels in a 5x5 pixel box around the kinetochore were summed together over all planes. The same operation was performed at areas outside the spindle and subtracted from the kinetochore intensity. Time points with no detectable CenpC were not analyzed.

To calculate tubulin intensity around a given kinetochore (Figs. 3 and 4), two intensity linescans (1.5 µm long) were taken perpendicular to the sister kinetochore axis: one positioned 0.7 µm towards the centromere (Tub_in_) and one 0.7 µm away from the centromere (Tub_out_). To synchronize traces to the beginning of Mad1 loss, traces were examined visually to locate the time where Mad1 levels dropped while CenpC levels stayed constant, and t=0 was set for the time point immediately before such Mad1 loss began. The Mad1-to-CenpC ratio was then normalized to the average ratio from −100 s to 0 s. In Figs. 3E and 3H (no Mad1 loss), intensities were normalized to K2’s Mad1 intensity in the first 100 s and 300 s of the trace, respectively.

**Other measurements:** Kinetochore-to-pole distances (Fig. 1D) were calculated by projecting where an individual kinetochore fell on the pole-to-pole axis. The “near” pole was designated as the pole closest to K1 at the time of K1’s Mad1 loss start.

**Statistics:** Data are expressed as mean ± SEM. Calculations of p-values (Student’s t-test) were done in StatPlus.

## Online supplementary material

Figure S1 describes the spatiotemporal trajectory of sister kinetochores around Mad1 loss: the time delay between sisters losing Mad1, the motion and position of sisters as they lose Mad1, and the orientation of the sister pair relative to the spindle around Mad1 loss. Figure S2 shows images of Mad1 loss in monopolar cells and Hec1-9A cells, and quantifies K-K distance in these cells. Video 1 shows an example trajectory of Mad1 loss on both kinetochores in a pair. Video 2 shows an example of Mad1 loss in amonopolar spindle. Video 3 provides an up-close view of a kinetochore pair as it loses Mad1 in order to illustrate centromere tension changes. Video 4 illustrates lateral microtubule attachment at a Mad1-positive kinetochore under persistent tension. Video 5 demonstrates that Mad1 levels are stable at CenpE-rigored (GSK-923295 treated) laterally attached kinetochores that remain stuck away from the metaphase plate. Video 6 shows how Mad1 leaves rapidly after end-on microtubule attachment begins.

## Acknowledgements

We thank Jagesh Shah for EYFP-Mad1 PtK2 cells, Michael Davidson for the mCherry-α-tubulin and mRuby2 constructs, Aaron Straight for the mCherry-CenpC construct, Jennifer DeLuca for the Hec1-9A-EGFP construct, Xiaohu Wan for Matlab SpeckleTracker, Kurt Thorn for microscopy and image analysis advice, and David Morgan, Fred Chang, and the Dumont lab for discussions and critical reading of the manuscript. This work was funded by NIH DP2GM119177 (S. D.), the Rita Allen Foundation and Searle Scholars’ Program (S. D.), and a NSF GRF (J. K.). The authors declare no competing financial interests.

## Author contributions

J. K. performed experiments and analyzed the data. J. K. and S. D. conceived the project, designed experiments and wrote the manuscript.

## Abbreviations list

SAC (spindle assembly checkpoint); k-fiber (kinetochore-fiber); STLC (S-trityl-L-cysteine); K1/2 (first/second sister kinetochore in a pair to lose Mad1)

